# Beyond Conveyance: Assessing the Contribution of Process-Critical Bacteria from Urban Sewer Systems to Wastewater Treatment Plants

**DOI:** 10.64898/2026.09.11.749817

**Authors:** Rodrigo Maia Valença

## Abstract

The sewer microbiome is known to be the main source of process-critical bacteria involved in nitrogen and phosphorus removal in wastewater treatment plants (WWTPs) and is thus crucial for the assembly of activated sludge (AS) microbial communities. However, the extent of these bacterial loads arriving at WWTPs currently lacks scientific attention. We provide a quantitative description of the biomass loads from sewer systems to WWTPs by coupling microbial community data by 16S rRNA amplicon sequencing from the Aalborg municipality sewer catchment with mass balances and biomass production across gravity and pressure sewer systems. Accounting for microbial transformations in sewer wastewater, biofilm and sediments under steady state dry weather flow conditions, gravity sewers were found to produce higher biomass loads than pressure systems, with loads of 1.57 and 0.27 kg_COD_ m^−3^ d^−1^ respectively. A key feature of our sewer model is its ability to estimate biomass loads from specific taxa, such as polyphosphate accumulating organisms (PAOs). Using Aalborg West WWTP as a case study, model estimations correlated well with figures calculated based on influent wastewater data monitoring. The model-estimated loads for the PAOs *Azonexus* and *Ca*. Accumulibacter were 2.10 kg day^−1^ and 1.80 kg day^−1^, compared to the influent wastewater monitoring-based loads of 1.77 kg day^−1^ and 1.20 kg day^−1^, respectively. While providing a framework for further research on sewer microbial dynamics, this study offers a tool for gaining insights into microbial immigration, generating valuable information for improving microbial communities management in WWTPs, optimizing nutrient removal, and enhancing WWTP design.

**Graphical abstract:** 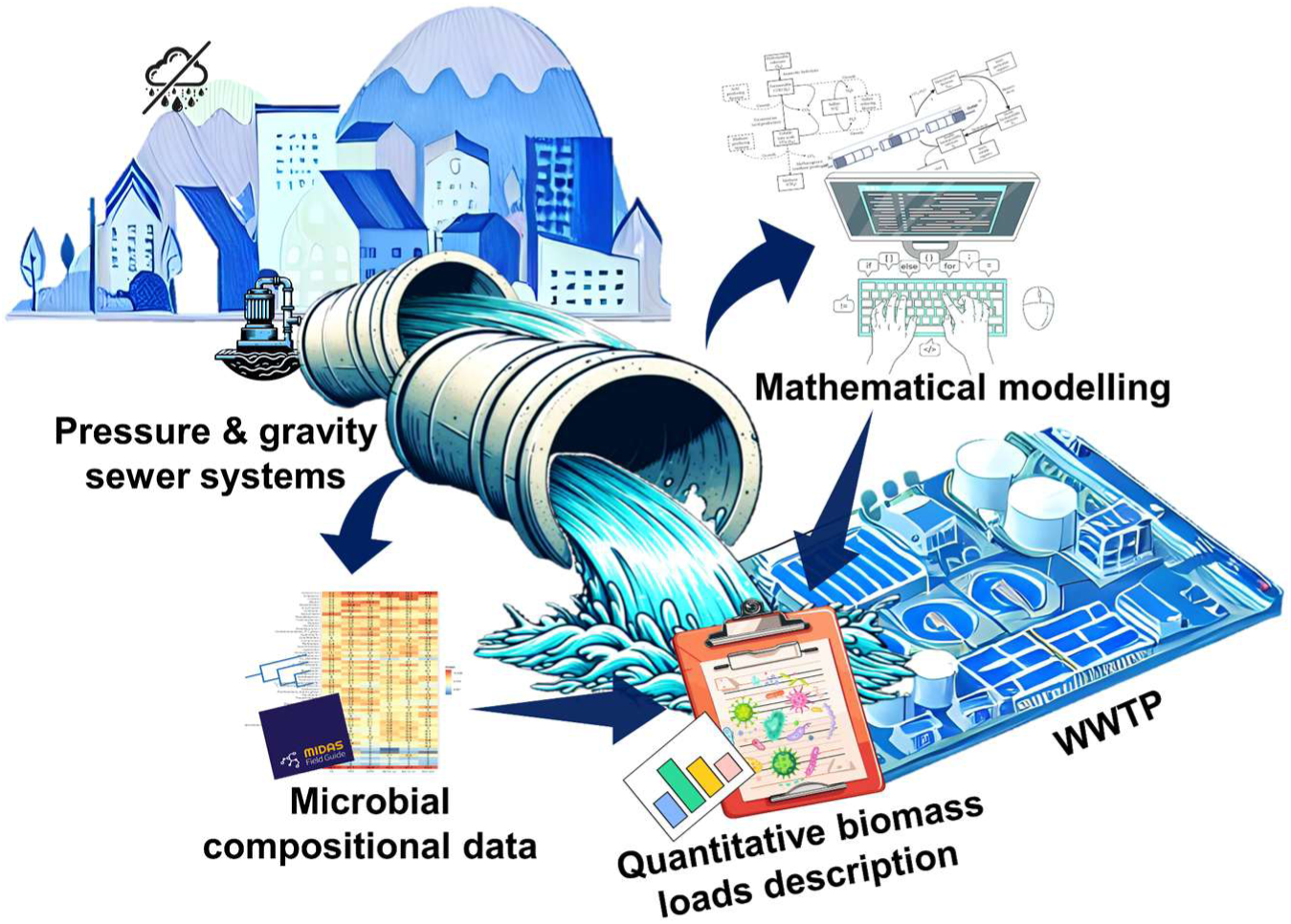

**Highlights:**

❖ Sewer biomass loads were estimated considering wastewater, biofilm and sediment
❖ Mean biomass production in gravity and pressure sewers is 1.57 and 0.27 kg_COD_ m^−3^ d^−1^
❖ Sewer provides ~3000 kg d^−1^ of biomass to the WWTP, consistent with model estimations
❖ A novel feature of the sewer model is estimating biomass loads from specific taxa
❖ *Azonexus* and *Ca*. Accumulibacter are more abundant in pressure than gravity sewers

## 1. Introduction

Activated sludge (AS) is among the most commonly used processes in wastewater treatment plants (WWTPs), with microorganisms playing a crucial role in pollutant removal and nutrient recovery. Despite the significant efforts made in recent years to characterize AS microbial communities (Wu et al., 2019; Dueholm et al., 2022), many factors influencing their microbial composition remain poorly understood. WWTPs are open ecosystems, where the AS bioreactor receives influent wastewater (IWW) containing both pollutants and microorganisms. Recent studies have shown that the community composition of AS is strongly influenced by the microbial communities present in the influent wastewater (Dottorini et al., 2021; Sun et al., 2021; Gibson et al., 2024), so there is an increasing interest in understanding the composition and sources of the incoming microorganisms.

The microbial communities present in influent wastewater reach WWTPs via sewer systems. As critical components of urban infrastructure, sewers are designed to protect cities from flooding and prevent the spread of waterborne diseases by conveying wastewater to WWTPs (Butler et al., 2018). They also provide an environment with plenty of substrate availability for microbial growth (Hvitved-Jacobsen et al., 2013), encompassing a large reservoir of biomass relevant to WWTP processes. Most studies concerning sewer microbial communities have focused on specific functional groups relevant to sewer management, such as sulfate-reducing bacteria linked to concrete corrosion (Cayford et al., 2017; Yuan et al., 2022) and methane producers, a greenhouse gas that poses safety risks in sewers due to its explosive properties (Song et al., 2023). Few studies have assessed overall microbial communities in sewers, and those that have were based on samples collected from influent wastewater at WWTPs (McLellan et al., 2019; LaMartina et al., 2021; Roguet et al., 2022).

In a recent study, we described microbial communities based on samples taken directly from sewer systems upstream of WWTPs (Riisgaard-Jensen et al., 2025). We found an extensive sewer microbiome, i.e., bacteria that were growing in the sewer sediment and biofilms, and, interestingly, we identified many of these to be process-critical bacteria in the AS-WWTPs. Our findings also underscored differences in microbial communities of sewer biofilms, sediments, and wastewater, and differences among gravity and pressure systems communities.

The driving force behind wastewater movement in sewer networks determines whether aerobic or anaerobic conditions prevail within the system. Potential energy differentials govern wastewater flow in gravity sewers, which are designed to operate partially full. This design ensures open channel conditions and ventilation, favoring aerobic processes to dominate in the wastewater and the outer layers of biofilms and sediments. In contrast, pressure sewers involve intermittent pumping of wastewater in fully flowing pipes, inducing anaerobic conditions (Hvitved-Jacobsen et al., 2013). In general, aerobic bacteria grow faster than anaerobic bacteria. Additionally, the hydraulic conditions in pressure sewers impose greater shear stress on biofilms than those in gravity sewers, leading to generally thinner biofilms. Consequently, the dynamics of gravity and pressure sewers are expected to produce different amounts of biomass, underscoring the importance of separately assessing biomass loads from each system.

The influence of sewer biomass on WWTP processes has long been acknowledged (Warith et al., 1998). However, despite this recognition, it remains unclear how much biomass is transported from sewer systems to WWTPs. Direct quantification of in-sewer biomass is challenging due to the dynamic nature, extensive length, and difficult access of sewer systems (De Feo et al., 2014). As a result, modeling approaches have been employed to enhance our understanding of sewer processes and management. These models, however, have primarily focused on organic matter and pollutant transformation, or gas production and sewer corrosion (Hvitved-Jacobsen et al., 1998; Jiang et al., 2009; Sun et al., 2018), with little attention given to sewer biomass and microbial ecology. A well known example is the Wastewater Aerobic/Anaerobic Transformations in Sewers (WATS) model, a conceptual sewer process model that has been used for analyzing concrete corrosion and odor nuisances caused by hydrogen sulfide, as well as several other wastewater compounds transformation (Hvitved-Jacobsen et al., 2013).

In this study, we uncover the differences in microbial compositions between gravity and pressure sewer systems, and provide a quantitative description of biomass loads from sewer systems under dry weather conditions by linking mathematical modeling – based on suspended biological growth and biofilm detachment – with sewer microbial compositional data. This approach enables the assessment of not only the overall sewer biomass loads but also the loads of specific functional groups arriving at WWTPs, such as polyphosphate accumulating organisms (PAOs) and glycogen accumulating organisms (GAOs) critical for AS performance. This information provides better understanding of factors controlling microbial community assembly, contributing to better community management, nutrient removal, and enhanced WWTPs design, stressing the strong link between sewer systems and WWTP performance.

## 2. Materials and methods

### 2.1. Model framework

The simulation of microbial biomass production in sewer systems conveyed to the wastewater treatment plant was based on the WATS model (Hvitved-Jacobsen et al., 2013). The WATS model was adapted to incorporate biofilm growth and detachment to determine the total in-sewer biomass production (Tables S1-S5). The proposed model simulates sewer microbial transformations based on the sewer infrastructure of the Aalborg municipality, Denmark, under average daily dry weather flow conditions and steady-state conditions. As an overview, the Aalborg municipality sewer infrastructure consists of 70% separated sewers and 30% combined sewers, spanning over 2000 km of sewer pipelines in total. Of these, around 90% of the length are gravity sewers and 10% are pressure mains, with average pipe diameters of 250 mm and 140 mm for gravity and pressure sewer systems, respectively. Since the sewer type controls sewer biochemical processes, both gravity and pressure sewers were considered and modeled according to European standard design criteria (Linde et al., 2011; Blazejewski et al., 2012). Gravity sewers were designed to ensure self-cleaning velocities of 0.6 m s^−1^ with wastewater flowing at 10% of pipe capacity in combined sewers and 35% in separated sewers (Butler et al., 2018). Pressure sewers had an in-network retention time of 6 hours in 1.5 km long, full-flowing pipelines, similar to conditions reported in the literature (Sun et al., 2014; Sun et al., 2018).

The main difference imposed by the sewer types on the microbial communities is the availability of oxygen as the final electron acceptor. In partially filled gravity sewers, reaeration ensures the predominance of aerobic conditions, favoring hydrolysis and aerobic organic matter oxidation, therefore all biomass in gravity systems was assumed to be aerobic heterotrophs (Hvitved-Jacobsen et al., 1998). In contrast, anaerobic conditions are predominant in full flowing pressure systems, where microbial processes include hydrolysis, acidogenic fermentation, acetogenesis, sulfidogenesis, and methanogenesis. In pressure systems, the microbial biomass was assumed to be composed of fermentative microorganisms in the bulk water and acetogenic, methanogenic, and sulfidogenic microorganisms in biofilms (Hvitved-Jacobsen et al., 2013; Sun et al., 2014). Also, to best model the sewer processes, wastewater organic matter was fractionated into readily biodegradable substrate, fast hydrolyzable substrate, slow hydrolyzable substrate, and biomass (Hvitved-Jacobsen et al., 2013). Sewer biofilms were assumed to be distributed along the entire pipe wetted perimeter to also account for sediment deposition, and assumed steady-state conditions cause sewer biofilms to maintain constant thickness due to the equilibrium between growth and detachment rates. Therefore, biofilm detachment equals biofilm growth (Hvitved-Jacobsen et al., 2013; Li et al., 2019), allowing biofilm growth to be modeled using Monod-type kinetics (Horn et al., 2003; Rittmann and McCarty, 2020) and accounted for as biofilm detachment, as seen in Tables S2-S5.

The WATS model employed in the current study was not calibrated. Instead, the model was set up and run with typical model parameters determined in previous studies based on similar systems (see Table S1 in the Supplementary Information). For evaluating the impact of model parameter selection on the simulation results, a sensitivity analysis was performed (Section 3.2). The modeled process rates and model matrices for gravity and pressure sewers are listed in Tables S2-S3 and Tables S4-S5, respectively. Reaeration in gravity sewers was calculated using Equation S1, and the effect of temperature on reaeration and reaction rates was calculated according to Equation S2. The model was implemented in R (https://www.R-project.org/), and the script is available at https://github.com/rmvalenca/SewerBiomass.

### 2.2. Sewer biomass production key metrics

The main model outputs are the biomass concentration in the water phase and the portion of biomass attributed to biofilm growth and detachment. The total biomass production over sewer networks consists of three components: the initial amount of biomass entering the system within the conveyed wastewater, the biomass growth in the bulk water phase, and the biomass from detached biofilm. Two key metrics are derived from the model: biomass production rate (BPR) and biofilm detachment rate (DR). These metrics serve as reference values that can be applied across various sewer systems, assisting the estimation of biomass loads from different systems with minimal input required. BPR and DR are calculated as described in Equations 1 and 2.

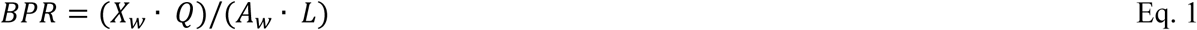

where:

*BPR* is the daily biomass production per wastewater volume (kg_COD_ m^−3^day^−1^);
*X_w_* is the biomass concentration in the water phase (g_COD_ m^−3^);
*Q* is the wastewater flow (m^3^day^−1^);
*A_w_* is the cross-sectional wetted area (m^2^);
*L* is the pipe length (m).

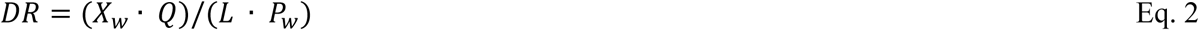

where:

*DR* is the daily biofilm detachment per pipe surface area (g_COD_ m^−2^day^−1^);
*X_w_* is the biomass concentration in the water phase (g_COD_ m^−3^);
*Q* is the wastewater flow (m^3^day^−1^);
*L* is the pipe length (m);
*P_w_* is the wetted perimeter (m).

### 2.3. Aalborg Sewers catchment and Aalborg West WWTP data

Information regarding the Aalborg municipality sewer catchment was obtained from the Aalborg Utilities Environment and Working Environment Report 2020 for the Water Division, (*Miljø-og arbejdsmiljøredegørelse 2020 for vanddivisionen)* (Aalborg Forsyning, 2021), and the Aalborg municipality wastewater management plan for 2021 to 2032 (*Spildevandsplan 2021-2032*) (Aalborg Kommune, 2021). Data on the physicochemical parameters of the Aalborg West municipal wastewater treatment facility (AAW) from 2018 to 2020 was provided by the plant operators and is available in Supplementary file 1.

### 2.4. Microbial data

This study was based on 74 sewer wastewater and biofilm grab samples collected between 2021 and 2023 from gravity sewer manholes, sewer overflow stations, and end of pressure sewers manholes within the Aalborg municipality sewer catchment, described in Riisgard-Jensen et al. (2025). In addition to 116 24-hour flow-proportional influent wastewater samples collected before primary settling, and 88 AS grab samples from AAW between 2019 and 2022, described in Riisgard-Jensen et al. (2023). All samples had their DNA extracted and amplicon sequenced using primers targeting the V1-V3 region of 16S rRNA genes as described in Dottorini et al. (2021). The sequencing was performed using Illumina MiSeq, and the data were processed using the AmpProc 5.0 workflow (https://github.com/eyashiro/AmpProc). Then, the data were mapped to the full-length amplicon sequence variants with the MIDAS 5.3 database (Dueholm et al., 2024). Samples with less than 5000 reads were removed and duplicate samples were merged by taking the mean read count. The data was processed, and heatmaps were generated using R v4.1.0 (https://www.R-project.org/) and RStudio 2022-02.0+443 (https://www.rstudio.com/), employing the packages ampvis2 (Andersen et al., 2018) and ggplot2 (Wickham, 2009). Scripts are available at https://github.com/rmvalenca/SewerBiomass and 16S data are available at the National Center for Biotechnology Information under Bioprojects PRJNA946374 and PRJNA1139651.

## 3. Results and discussion

### 3.1. Estimating Biomass Production in Sewer Systems

A model framework was implemented to quantify the total biomass production in sewers and evaluate its significance on downstream WWTPs. The biomass production was estimated by adapting the WATS model (Hvitved-Jacobsen et al., 2013) and using the sewer infrastructure of Aalborg municipality, Denmark, as a case study. Aalborg municipality sewer catchment area comprises over 2000 km of sewer pipelines, with approximately 70% separated sewers and 30% combined sewers. Of these, 90% are gravity sewers and 10% are pressure mains. The average pipe diameter is 250 mm for gravity sewers and 140 mm for pressure mains transporting average dry weather flow of 1.5 L s^−1^ with a total organic matter concentration of 600 g_COD_ m^−3^. Based on previous reports, sewer wastewater temperature was set to an average of 10 °C (Kretschmer et al., 2016; Johra and Heiselberg, 2018), and the dissolved oxygen in gravity sewer wastewater was assumed to be 4 g_O2_ m^−3^ (Hvitved-Jacobsen et al., 2000). The average biofilm thickness, based on previous literature, was assumed of 1 mm in gravity sewers (Hvitved-Jacobsen et al., 2013; Li et al., 2019) and 0.6 mm in pressure sewers (Hvitved-Jacobsen et al., 2013; Sun et al., 2014), distributed along the entire wetted perimeter to account for sediment deposition. And, on average, biofilms were assumed to contain 10 g_COD_ m^−2^ of active biomass (Hvitved-Jacobsen et al., 2000).

The referred model can be used to estimate the sewer biomass loads over a known sewer catchment, here expressed as the biofilm detachment rate (DR) and biomass production rate (BPR) (Table 1). The sewer type has a strong impact on the biofilm production rate (equal to DR). The active microbial communities in gravity sewers are predominantly aerobic (Hvitved-Jacobsen et al., 2013), comprising faster-growing bacteria compared to the anaerobic bacteria typically found in pressure systems. The faster microbial growth rates of heterotrophic aerobes as well as differences in biofilm density, maturity level, and hydraulic stress (Hvitved-Jacobsen et al., 2013) cause gravity biofilms to grow and detach faster than pressure biofilms. Assuming there are 10 g_COD_ m^−2^ of active biomass in sewer biofilms, the calculated DRs for average sewer sizes and 1 km length were 1.31 g_COD_ m^−2^ day^−1^ for gravity biofilms and 0.09 g_COD_ m^−2^ day^−1^ for pressure biofilms (Table 1). The big difference also results in significant differences in time for renewal of the biomass, less than 10 days for gravity biofilms and around 100 days for biofilms in pressure systems. Similarly, when considering the biomass growth in the bulk water phase, gravity sewers produce higher BPR than pressure systems (1.57 and 0.27 kg_COD_ m^−3^ day^−1^, respectively). This highlights the difference in microbial growth dynamics between the two systems.

**Table 1.** Standing sewer biomass both on biofilm and bulk water phase, biomass production rate (BPR), and biofilm detachment rate (DR) of the Aalborg municipality sewer systems. Based on one kilometer of steady-state sewer microbial transformations, considering 10% full gravity and full flowing pressure sewers, with biofilm thicknesses of 1 mm and 0.6 mm, transporting wastewater flow of 1.5 L s^−1^ with 600 mgCOD L^−1^ at 10 °C.

|  | Gravity sewers | Pressure sewers |
| --- | --- | --- |
| Active biomass in biofilm (g<sub>COD</sub> m<sup>-2</sup>)* | 10 | 10 |
| Active biomass in bulk water (g<sub>COD</sub> m<sup>-2</sup>)** | 0.48 | 1.10 |
| DR (g<sub>COD</sub> m<sup>-2</sup> day<sup>-1</sup>) | 1.31 | 0.09 |
| BPR (kg<sub>COD</sub> m<sup>-3</sup> day<sup>-1</sup>) | 1.57 | 0.27 |

\*Average estimate from literature Hvitved-Jacobsen et al. (2000).
\*\*Calculated considering BPR and DR, with 250 mm diameter gravity sewers, and 140 mm pressure sewers.

Further focusing on the Aalborg municipality sewer catchment, based on the amounts of active biomass in sewer biofilm and bulk water (Table 1), it can be estimated that the entire sewer catchment holds ~0.2 tons_COD_ of standing biomass (defined as the amount of biomass occupying an area at a particular time) in sewer bulk water and ~2.8 tons_COD_ in sewer biofilms distributed among sewer networks connected to the two wastewater treatment facilities. The largest one, Aalborg West WWTP (AAW), processes wastewater from approximately 55% of the municipality’s sewer catchment, covering around 1000 km of pipelines. AAW receives average dry weather flows of 45,000 m^−3^ day^−1^ of wastewater with organic matter concentration of 600 (407 −870) g_COD_ m^−3^.

Based on this description and considering that 10% of the influent organic matter, measured as COD, corresponds to total cell biomass (Foladori et al., 2010), we used our model to simulate AAW sewer catchment, estimate the biomass loads from its sewer pipelines, and compare that to the AAW influent biomass loads calculated from the plant’s monitoring data (Supplementary File 1). The biomass loads from the model estimation of 2.93 tons_COD_ per day agrees well with the biomass loads from the AAW monitoring data of 2.70 (range 1.83 −3.91) tons_COD_ per day, validating the modeled processes and assumptions. The total AAW influent daily biomass loads (calculated from Supplementary File 1 data) correspond to 13% (range 12% - 15%) of its AS biomass, highlighting the role of the sewer systems input for the community assembly in the WWTPs.

### 3.2. Impact of Sewer Parameters on Sewer Biomass Production

The size and functionality of a sewer network impact its microbial processes in different ways. For example, small-diameter pressure pipes have a larger surface area-to-volume (A/V) ratio, which reflects the distribution of standing biomass. This higher A/V ratio means there is significantly more biomass in biofilm form (surface-related grown bacteria) than in the bulk water phase (suspended-related grown bacteria), leading to increased odor potential and gas emissions compared to larger-diameter pipes (Hvitved-Jacobsen et al., 2013). Furthermore, urban sewer systems undergo a series of changes before reaching their final destination. For example, sewer pipe diameters can vary to accommodate different catchment characteristics and configurations, ranging from 100 to 1000 mm in the Aalborg municipality sewer catchment. Additionally, the daily average organic matter concentration in sewer wastewater varies with weather conditions and overall water consumption; at the Aalborg West WWTP, influent wastewater organic matter ranges from 240 to 1300 g_COD_ m^−3^ over the year. Sewer wastewater temperature was assumed to be the same as the Danish soil temperature 2 meters below ground, then the yearly sewer temperature variance would be 5 to 15°C (Johra and Heiselberg, 2018). Solids tend to accumulate in sewer pipes, and sewer biofilm thickness varies according to sewer configuration, biofilm development, and hydraulic conditions. Pressure sewer biofilms typically range between 0.3 and 0.9 mm in thickness (Hvitved-Jacobsen et al., 2013; Sun et al., 2014), while gravity sewer biofilms (and sediments) can reach up to 2 mm (Li et al., 2019).

To better understand the magnitude of these parameters’ influence on the modeled sewer biomass production, we varied pipeline diameters from 70 to 280 mm for pressure sewers and from 125 to 500 mm for gravity sewers, biofilm thickness from 0.3 to 1.2 mm for pressure systems and from 0.5 to 2 mm for gravity systems, substrate concentration from 300 to 1200 g_COD_ m^−3^, and wastewater temperature from 5 to 20°C. The results from this sensitivity analysis show that both modeled sewer types respond very similarly to changes in these parameters (Figure 1) for DR and DPR responses for variations in pressure and gravity sewers. Both BPR and DR are independent of the diameter of the pipe when considering a constant flow depth of 10% of the pipe’s capacity for gravity sewers and fully flowing pressure sewers. Additionally, the bacteria were assumed to be evenly distributed across biofilm depth, causing biofilm thickness to affect the amount of reacting biomass in biofilms, linearly impacting BPR and DR. Substrate concentration also linearly affects sewer BPR in both gravity and pressure systems. However, the relationship between biofilm detachment and substrate concentration is only linear for pressure sewers (Figure 1a), where organic matter is the only substrate promoting microbial growth. In contrast, biofilm growth and detachment in gravity systems are limited by dissolved oxygen availability (Figure 1c). Furthermore, temperature exponentially affects biofilm production and detachment within sewer pipes due to the high temperature dependence of the process reaction rates (Equation S2).

**Figure 1.**
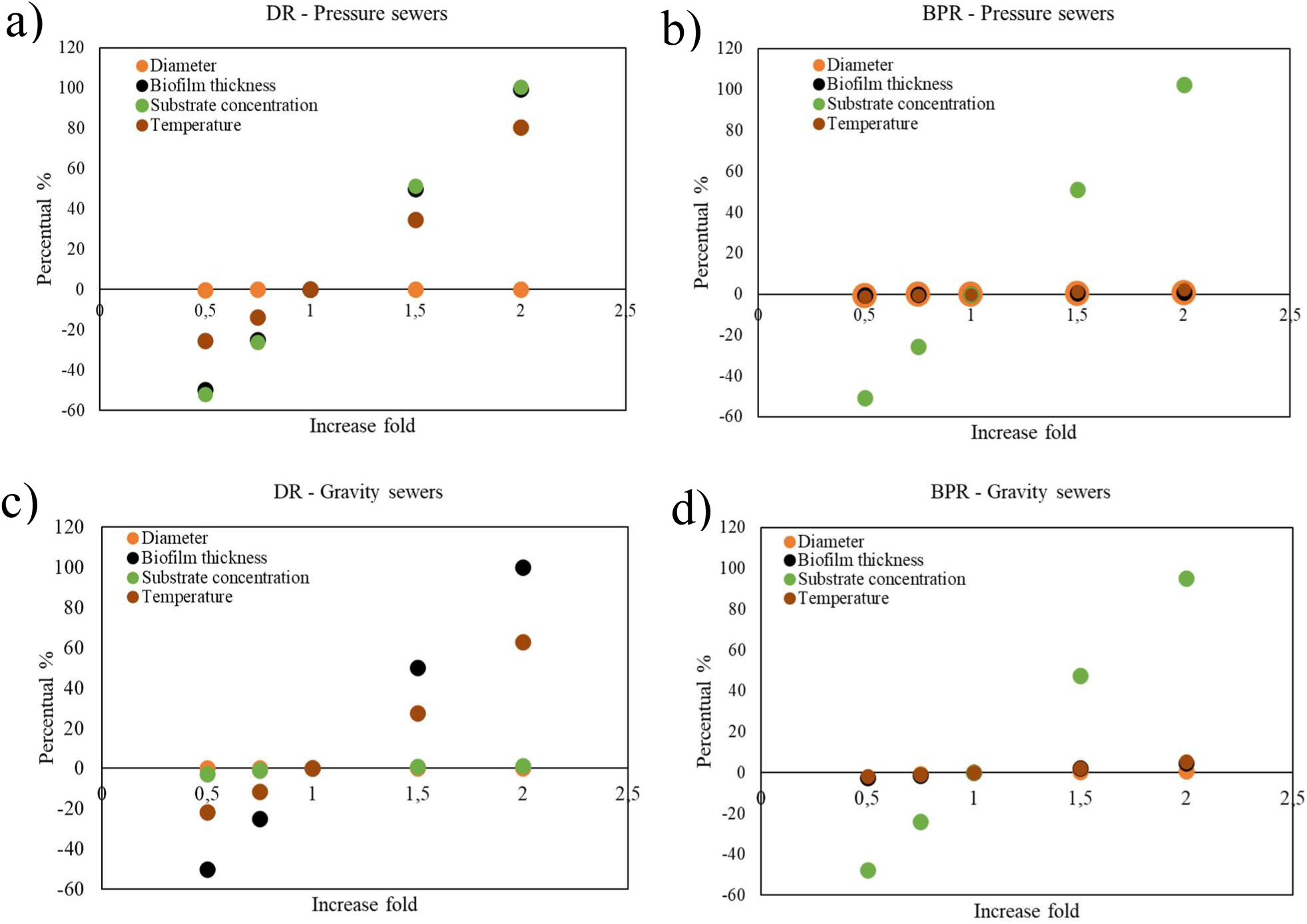
DR and BPR response to variations in wastewater substrate concentration, temperature, biofilm thickness, and pipeline diameter in different sewer types: a) DR - Pressure sewers, b) BPR - Pressure sewers, c) DR - Gravity sewers, and d) BPR - Gravity sewers.

Although BPR and DR are independent of pipe diameter, the total biomass production of sewer pipelines is linked to their diameter, because larger pipes transport greater wastewater volume, yielding greater biomass production. Additionally, the amount of biomass detached from biofilms is linked to the wetted perimeter. Therefore, larger diameter pipes, with their greater wetted perimeter and larger biofilm surface areas, lead to more biofilm detachment. Another factor influencing total sewer biomass production is the pipeline length, which affects the residence time of the system. Hence, to illustrate the effect of both pipe diameter and length on sewer biomass production, we have modeled and compared two pressure systems: one with 400 mm in diameter transporting 12.5 L s^−1^ and the other with 283 mm in diameter transporting 6.2 L s^−1^, over 1 and 2 km (Table 2).

**Table 2.** Biomass production over 400 and 283 mm pressure sewer pipelines. Considering full flowing pipes, with 0.6 mm thick biofilm, wastewater with 600 mg_COD_ L^−1^ at 10 °C.

| Ø<br>(mm) | Length<br>(km) | V<br>(m<sup>3</sup>) | Q<br>(L s<sup>-1</sup>) | BPR<br>(kg m<sup>-3</sup>d<sup>-1</sup>) | DR<br>(g m<sup>-2</sup>d<sup>-1</sup>) | Bulk water<br>biomass<br>(kg d<sup>-1</sup>) | Biofilm<br>detachment<br>(g d<sup>-1</sup>) | Total<br>biomass<br>(kg d<sup>-1</sup>) |
| --- | --- | --- | --- | --- | --- | --- | --- | --- |
| 400 | 1 | 126 | 12.51 | 0.27 | 0.09 | 34.13 | 114 | 34.24 |
| 283 | 1 | 63 | 12.51 | 0.27 | 0.09 | 17.06 | 81 | 17.14 |
| 283 | 2 | 126 | 6.25 | 0.27 | 0.09 | 34.13 | 161 | 34.29 |

The 400 mm pipe transports twice as much wastewater flow as the 283 mm pipe, resulting in double the biomass production over 1 km. To balance the flow difference, we have also considered the biomass production in the 283 mm pipe over 2 km, allowing both systems to transport the same wastewater volume, differing only in residence time. As a result, the total biomass production from wastewater transport differs by only around 0.1% between a 400 mm pipe over 1 km and a 283 mm pipe over 2 km, as shown in Table 2. However, the difference in the amount of biomass detached from biofilms is significant, at approximately 40%. Although the 283 mm pipe carries half the wastewater flow of the 400 mm pipe, the difference in wetted perimeters is not as substantial. Since biofilm detachment is caused by mechanical scouring from the flowing wastewater, the longer residence time in the 283 mm, 2 km long pipe results in the accumulation of more biomass from detached biofilm than the 400 mm, 1 km long pipe. While pipeline length and diameter are important factors influencing biomass production in similarly configured sewers, the most critical factor for total biomass production is the volume of wastewater conveyed. An important observation is that sewers with different combinations of length and diameter transporting the same wastewater volume will roughly produce the same amount of biomass.

### 3.3. The Impact of the Sewer Microbiome on the WWTPs

#### 3.3.1. Aalborg Municipality Sewers Microbial Overview

Assessing the relevance of sewer biomass loads to WWTPs requires consideration of both sewer biomass production rates and microbial composition. Figure 2 shows the most abundant bacterial genera in the Aalborg municipality sewers, divided into gravity sewer biofilms/sediments and water phase, pressure sewer biofilms and water phase, and the influent wastewater (IWW) to the AS plants.

**Figure 2.**
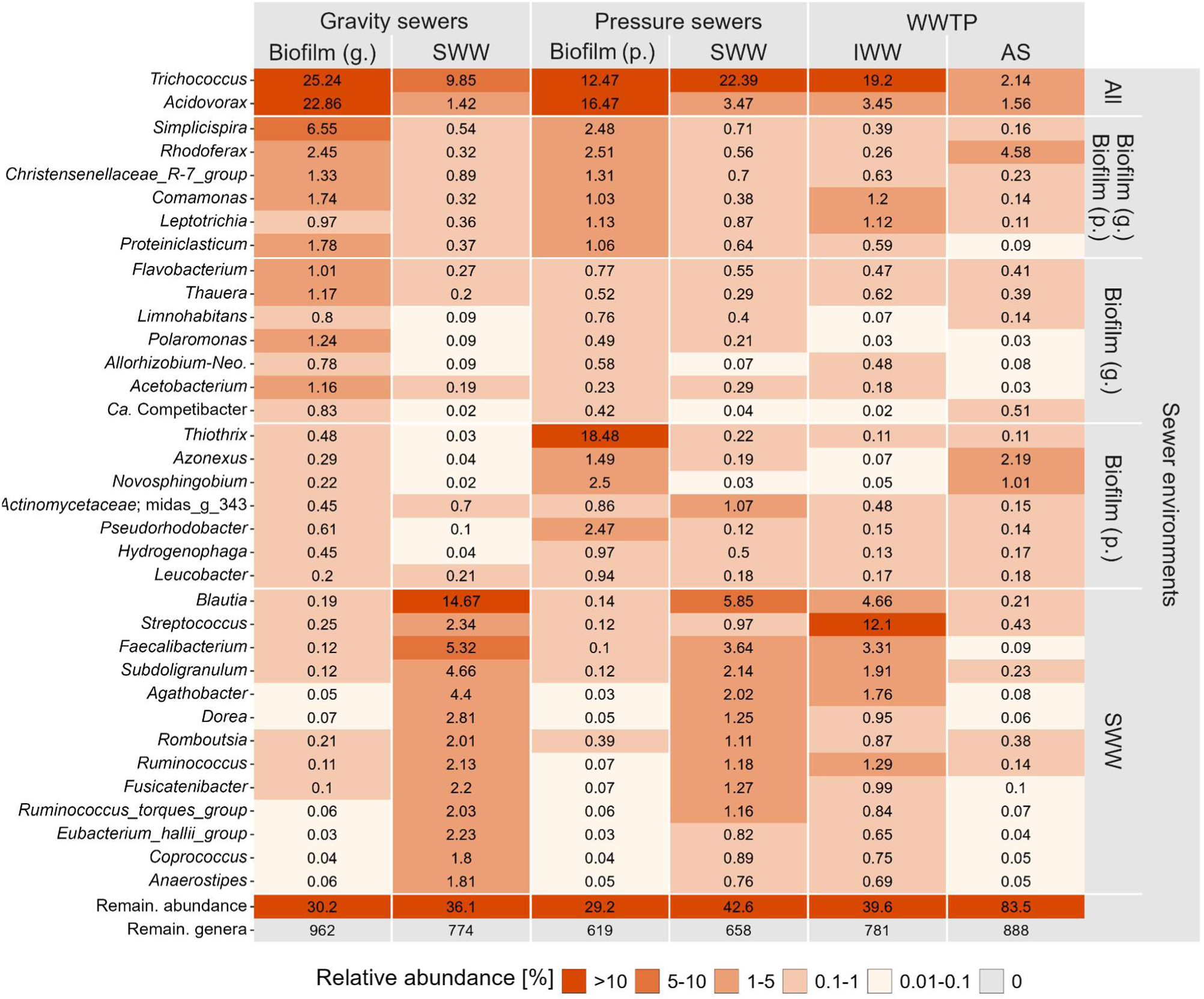
Top 15 most abundant genera among all sample types, biofilm (g.), biofilm (p.) and sewer wastewater (SWW). The top 15 SWW were found by combining pressure and gravity pipes. The relative abundance is the mean across all samples for each sample type. The facets indicate in which sample types the genera are part of the top 15. Abundances are shown with two decimals if ≥0.01. A relative abundance of 0 means no reads were detected. “Remain. abundance” and “Remain. genera” refers to the cumulative abundance or number of genera outside of the top 15, respectively. Only classified genera are included, with family names provided for genera classified with MiDAS placeholder names.

All abundant sewer microbes were also found in AS (Figure 2) where they either die off or grow as process-critical bacteria (Riisgaard-Jensen et al., 2025). Two genera, *Acidovorax* and *Trichococcus,* were consistently found across all sewer environments and they are part of the sewer microbiome, i.e., growing in the biofilms and/or sediments (Riisgaard-Jensen et al., 2025). *Acidovorax*, natural inhabitants of plants and soil, are primarily aerobic, but some can also grow anaerobically using nitrate as a terminal electron acceptor (Willems and Gillis et al., 2015; Siani et al., 2021). *Trichococcus*, fermentative organisms capable of both aerobic and anaerobic growth (Rainey, 2015), have been isolated from diverse natural environments such as guano, swamps, and high elevation wetlands (Strepis et al., 2020). The versatile metabolism of both genera make them omnipresent in sewers and WWTPs across the globe (Roguet et al., 2022; Riisgaard-Jensen et al., 2025).

Sewer wastewater comprises about 40% of gut bacteria (Riisgaard-Jensen et al., 2025) such as species within the *Dorea*, *Blautia*, and *Faecalibacterium* (Figure 2), making the sewer water phase more microbially diverse than biofilms. As wastewater continuously flows through the sewer pipelines, their microbial communities reflect the dynamic inputs from various wastewater sources. In contrast, biofilms develop and mature over time, naturally selecting organisms that best fit and adapt to their prevailing environmental conditions.

The sewer configuration significantly influences their microbial communities; partially full gravity sewers predominantly maintain aerobic conditions in the upper part of the biofilms/sediments, whereas full-flowing pressure systems foster anaerobic environments. One of the most notable disparities between the microbial compositions of gravity and end of pressure sewers is the relative abundance of *Thiothrix* in pressure sewer biofilms, comprising almost 20% of their bacterial community, as seen in Figure 2. The genus *Thiothrix* includes aerobic and facultative anaerobic, filamentous, non-motile, sulfur-oxidizing bacteria, typically found in flowing waters with an increasing gradient of sulfide and decreasing levels of oxygen (Ravin et al., 2022). This aligns with the environmental conditions at the locations from which the pressure sewer samples were taken. The availability of hydrogen sulfide from upstream anaerobic conditions, along with some level of oxygen exposure, might have favored the growth of *Thiothrix*, as previously described in the microbial community of corroding concrete in sewer systems (Okabe et al., 2007).

##### 3.3.1.1. Assessing top abundant bacterial loads from Aalborg municipality sewers to AAW

By combining the biomass production rates (Table 1) and the sewers’ microbial compositional data (Figure 2) through multiplication, it is possible to estimate the bacterial loads of the most abundant sewer taxa arriving at AAW and compare it to influent wastewater monitoring compositional data (Table 3).

**Table 3.**
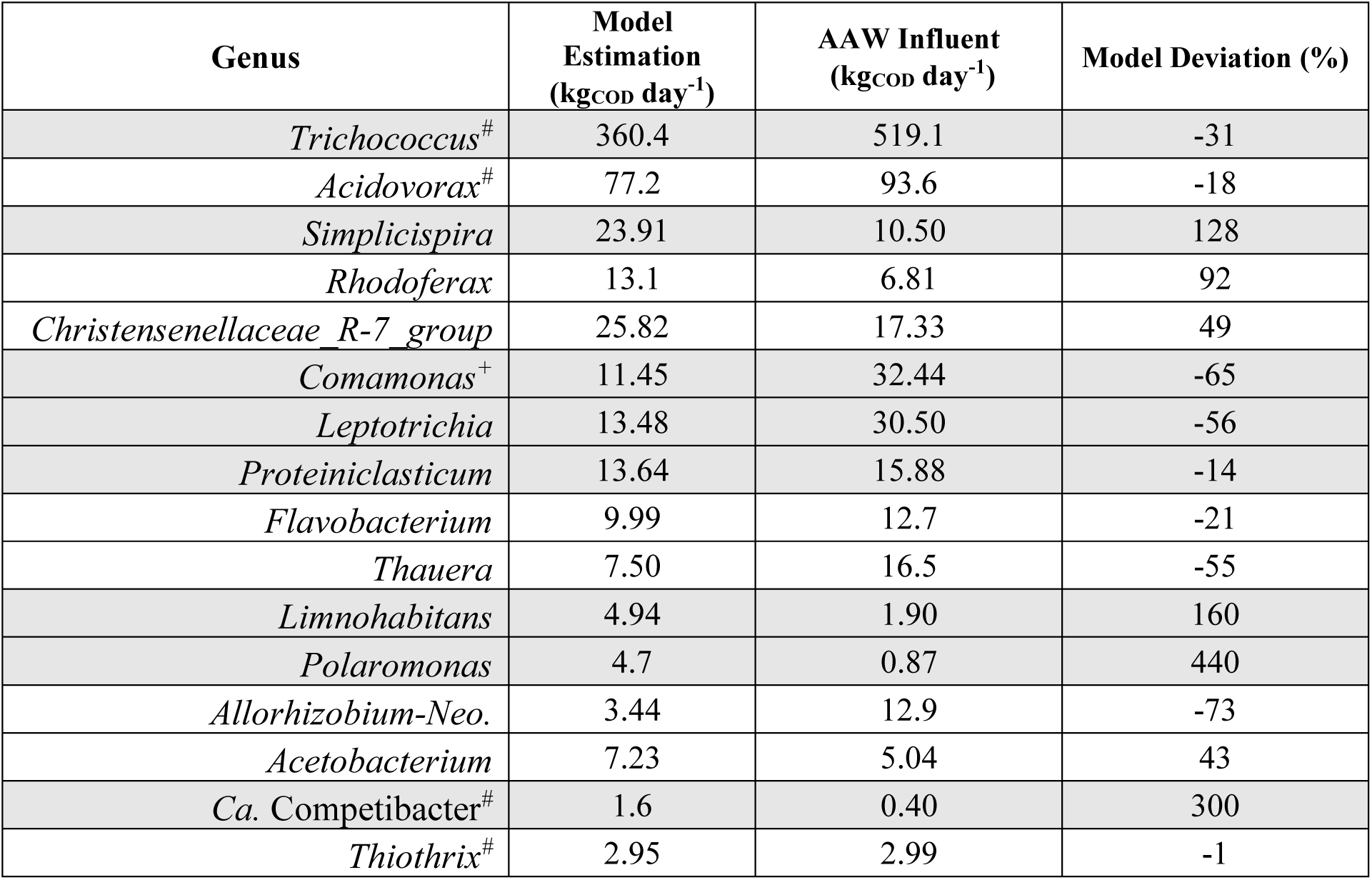
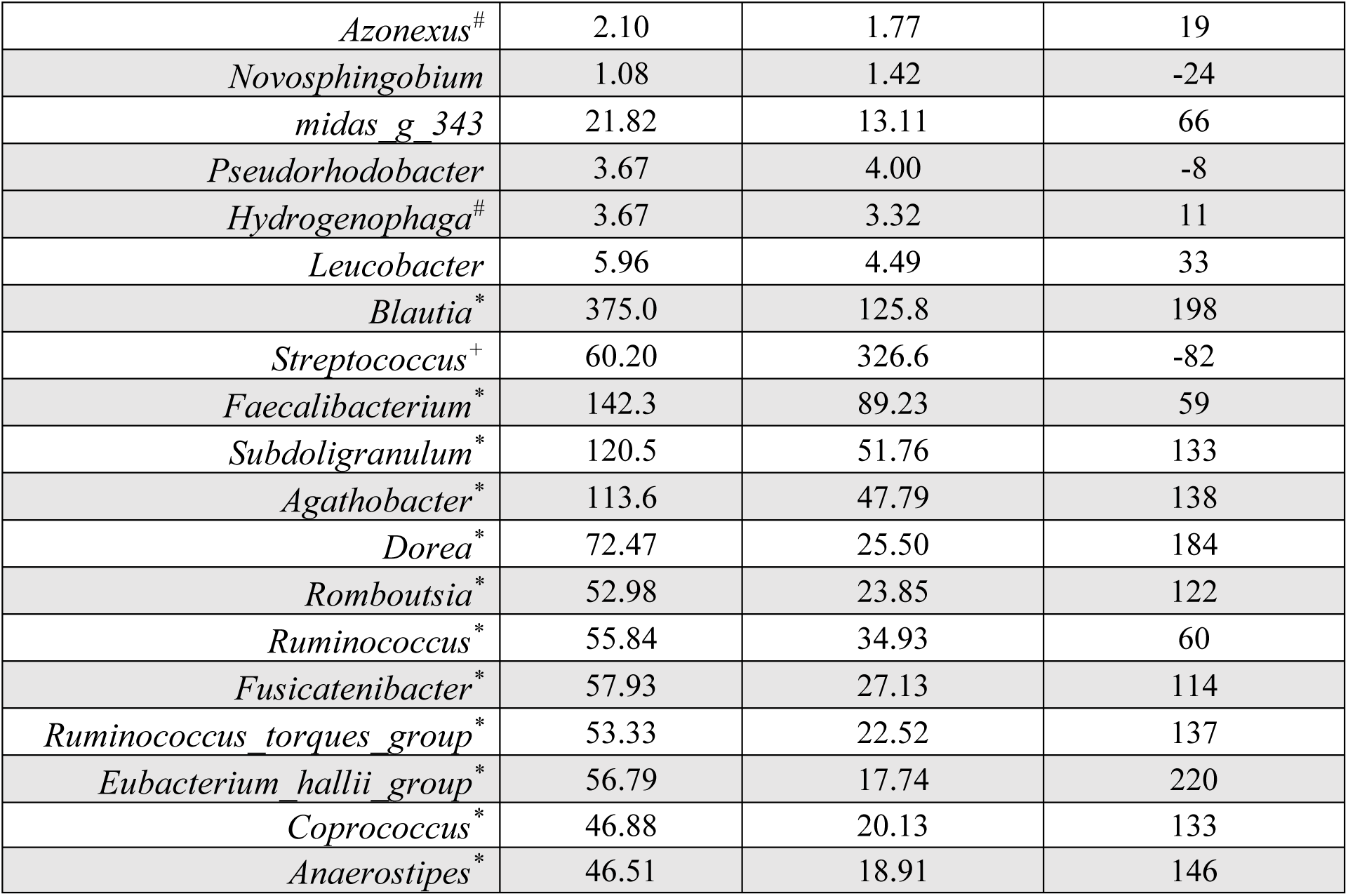
Comparison between the loads of the most abundant Aalborg sewer bacteria to the AAW, estimated by the proposed sewer model and from the WWTP monitoring data. Ordered by relative abundances in Figure 2. Model deviation defined as ((Model Estimated load - AAW Influent load)/AAW Influent load) × 100. *: Genera includes many human gut bacteria. +: Genera includes human pathogenic bacteria. #: Genera includes bacteria of the sewer microbiome (Riisgaard-Jensen et al., 2025).

The abundance of the bacterial genera in the influent wastewater (IWW) to AAW has been monitored for over one year and the load for the most abundant genera was between 0.4 to 519 kg_COD_ day^−1^. The model estimates showed overall the same level and trend for most genera. For example, it showed approximately ±30% precision for *Trichococcus* and *Acidovorax,* both belonging to the sewer microbiome. The results indicate that the sources of the microorganisms influenced the quality of the model’s estimates, with higher precision for organisms from the sewer microbiome, i.e. those growing in the sewer biofilms. Other bacteria, such as the gut bacteria *Blautia* and *Dorea* were almost 3 times overestimated by the model compared to AAW influent wastewater monitoring, while some human pathogens such as *Streptococcus* and *Comamonas* (Ryan et al., 2022; Timoney, 2022) were underestimated to around 30% of the AAW monitored load.

Bacteria not belonging to the sewer microbiome generally have poorly defined sources with the exception of gut bacteria, and may have specific sources or hotspots (e.g. hospitals) that have not been sampled and therefore not described well in the model. Other species, e.g. pathogens and gut bacteria are dying off during the transportation, a factor not included in the model. Gut bacteria are known to decay during sewer transportation (Riisgard-Jensen et al., 2025), so the difference in the estimated versus monitored abundances in the AAW influent wastewater might be attributed to the fact that sewer samples were taken very close to their sources, failing to provide enough time for the known gut bacteria to decay.

In a WWTP context, most of the high-abundance genera present in the sewers are known to decline in abundance in WWTPs, suggesting they are not relevant to the activated sludge process (Dottorini et al., 2021). Most of the AS process-critical bacteria responsible for C, N, and P removal are low-abundance genera in the influent wastewater to the WWTPs (Riisgard-Jensen et al., 2025). Therefore, to improve our understanding and capability to estimate the load of process-critical genera in WWTPs, potentially to optimize nutrient removal, we have chosen PAOs and GAOs as illustrative functional groups for a detailed study.

#### 3.3.2. Overview of PAOs and GAOs in Aalborg municipality sewers

AAW WWTP had a high diversity of PAO and GAO in the activated sludge with the most abundant genera being *Ca.* Phosphoribacter, *Azonexus* (previously named *Dechloromonas*), *Ca.* Accumulibacter, and *Tetrasphaera* for PAOs and *Ca.* Competibacter, *Defluviicoccus,* and *Propionivibrio* for GAOs (Figure 3). They were all present in IWW in both biofilms and sewer wastewater. The average compositional data values provide a satisfactory representation of the described systems, and the data variability across both sampling locations and sample types is illustrated in Figures S1 and S2. Overall, PAOs and GAOs genera were more abundant in sewer biofilms compared to wastewater, highlighting the observations of Riisgard-Jensen et al. (2025) that sewer biofilms and sediments serve as a source of these bacteria to WWTPs.

**Figure 3.**
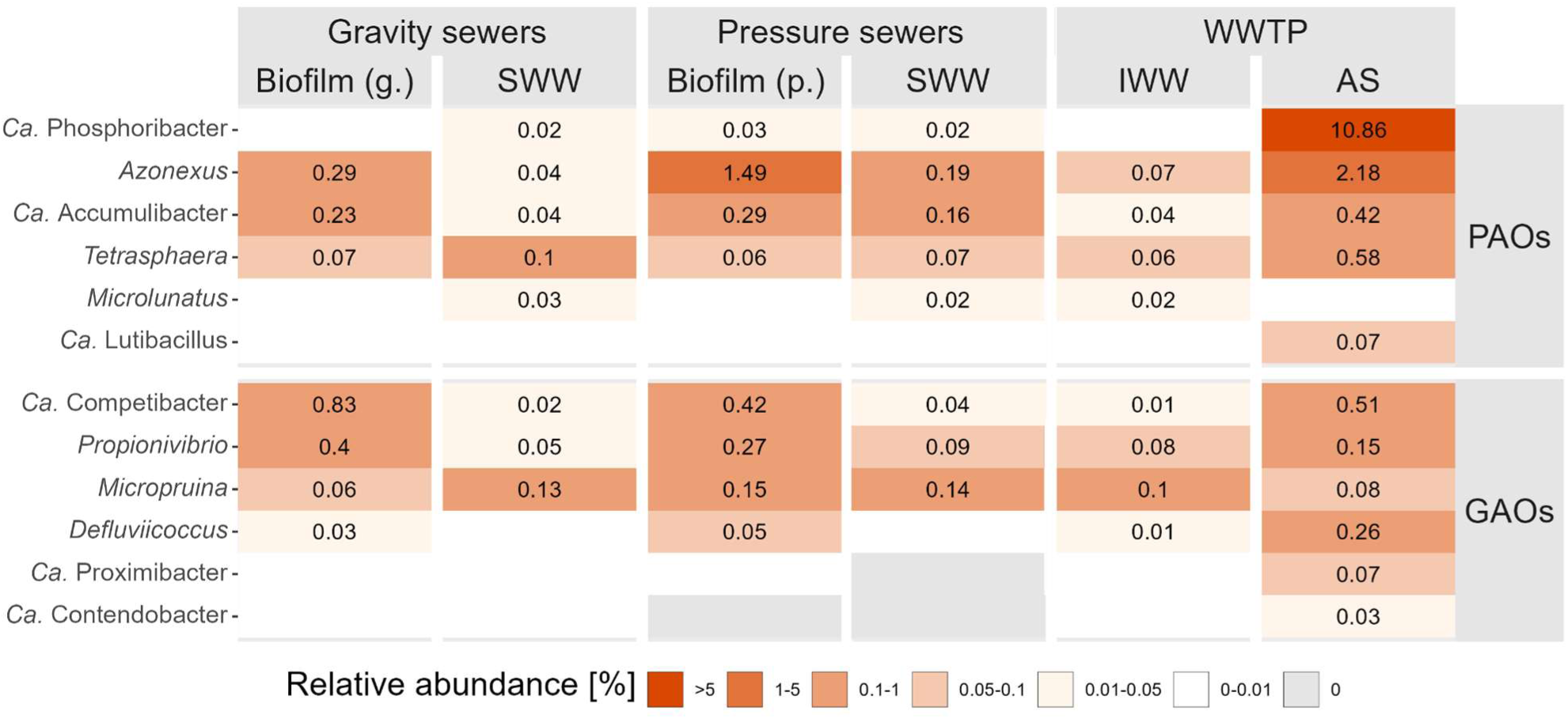
Relative abundance of polyphosphate accumulating bacteria (PAO) and glycogen accumulating bacteria (GAO) in Aalborg sewer catchment and Aalborg West WWTP. The relative abundances represent the average of all samples for each sample type (for sewer samples), and average of weekly samples for one year for IWW. The facets show whether the genera are a PAO or GAO. Abundances are shown with two decimals if ≥0.01. A relative abundance of 0 means no reads were detected.

The samples from pressure sewer locations were taken at the very end of each respective pressure line, where contact with atmospheric air made oxygen available. This, coupled with the typical accumulation of volatile fatty acids from pressure sewer microbial processes (Hvitved-Jacobsen et al., 2013), might have been responsible for the abundance of PAOs in the pressure sewer biofilms samples as they thrive in dynamic feast-famine conditions and with varying levels of oxygen. Aditionally, the presence of PAOs in gravity sewer biofilms also indicate that these biofilms undergo alternation of anoxic-oxic conditions, possibly due to variations in organic matter composition, and in wastewater flow depths, leading to fluctuations in the biofilm’s exposure to both wastewater and atmospheric air within the pipes (Saia et al.,2017).

*Azonexus* emerged as the most abundant PAO genus in both sewer configurations, reaching average relative abundance of 1.49% in pressure sewer biofilms. Additionally, *Ca*. Phosphoribacter, the most abundant PAO in Danish AS WWTPs (Dueholm et al., 2022), was only present in low abundance but were more abundant in pressure than gravity sewers, likely taking advantage of its fermentation capacity (Singleton et al., 2022).

##### 3.3.2.1. PAOs and GAOs loads from typical gravity and pressure sewers

Our modeling approach, combined with microbial compositional data (Figure 3), allowed us to estimate the PAO and GAO loads from average gravity and pressure sewers in the Aalborg sewer catchment (as described in section 3.1). To facilitate easy comparison between the two systems without the need for an illustrative example, the microbial loads in Table 4 were expressed per meter of sewer pipeline. Hence, the estimated PAO loads were 10.80 and 20.01 mg_COD_ m^−1^ day^−1^ for gravity and pressure sewers, respectively, while the GAO loads were 11.73 and 12.10 mg_COD_ m^−1^ day^−1^ for gravity and pressure sewers (Table 4). Although gravity sewers produce greater biomass loads, pressure sewers produce PAO-richer streams, highlighting the importance of accounting for the microbial composition of different sewer configurations when assessing sewer microbial dynamics.

**Table 4.**
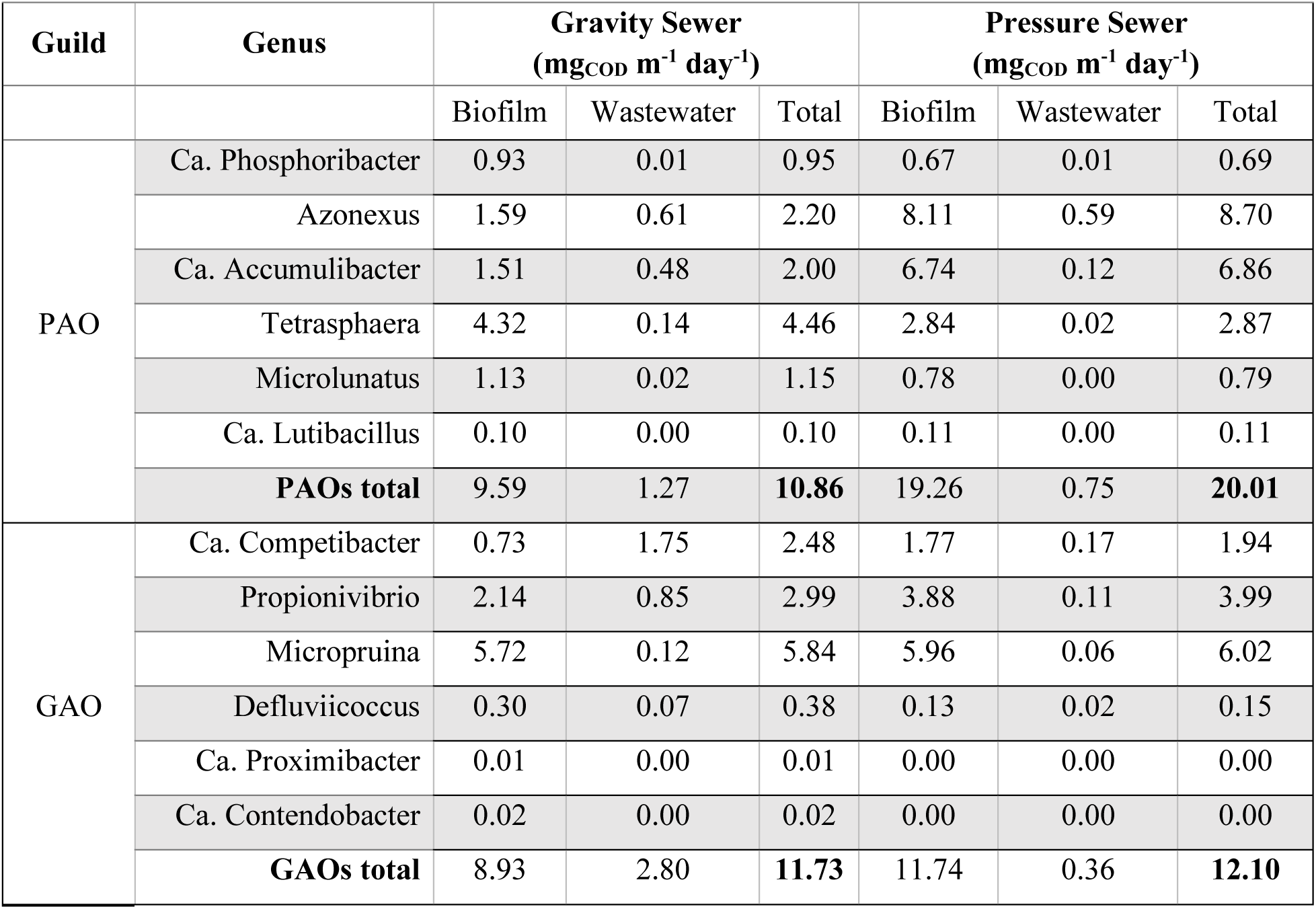
Typical PAOs and GAOs biomass loads from Aalborg municipality pressure and gravity sewers. Considering 10% full gravity and full flowing pressure sewers, with biofilm thicknesses of 1 mm and 0.6 mm, transporting wastewater flow of 1.5 L s^−1^ with 600 mg_COD_ L^−1^ at 10 °C.

##### 3.3.2.2. Enriching PAOs in WWTPs influent wastewater

Since PAOs play a crucial role in the P-removal performance of WWTPs, understanding their sources and finding ways to enhance their immigration is of significant interest. As pressure sewers show larger relative abundances of PAOs, a sewer catchment with a higher proportion of pressure mains compared to gravity sewers would tend to enrich PAOs in downstream WWTPs. In contrast, GAOs seem to be unaffected by sewer type. To assess how different sewer system configurations within a catchment influence PAOs and GAOs loads, we modeled and compared three configurations with varying gravity-to-pressure ratios: 100-0%, 50-50%, and 0-100% (Table 5).

**Table 5.** PAOs and GAOs biomass loads from the Aalborg sewer catchment had gravity-pressure sewer ratios of 100-0, 50-50, and 0-100%.

| **GRAVITY**<br>**SEWERS**<br>**(%)** | **PRESSURE**<br>**SEWERS**<br>**(%)** | **PAOS**<br>**(MG M<sup>-1</sup>DAY<sup>-1</sup>)** | **GAOS**<br>**(MG M<sup>-1</sup>DAY<sup>-1</sup>)** |
| --- | --- | --- | --- |
| **100** | **0** | 10.86 | 11.73 |
| **50** | **50** | 15.43 | 11.91 |
| **0** | **100** | 20.01 | 12.10 |

The total PAO loads nearly doubled, increasing from 10.86 to 20.01 mg m^−1^ day^−1^, when comparing only pressure sewers to only gravity sewers, while GAOs loads remained stable. Although it is not practical to change the amount of pressure mains in a sewer catchment solely to enrich PAOs at WWTPs, these findings illustrate the importance of sewer system configurations in shaping microbial composition of influent wastewater, which impacts on community assembly and WWTPs performance. This consideration may be valuable when planning and designing new sewer systems, not only for PAOs and GAOs but also for other process-critical bacteria, such as those involved in foaming and bulking.

##### 3.3.2.3. Assessing PAOs and GAOs loads from Aalborg municipality sewers to AAW

We compared the model estimates of the PAO and GAO load from the AAW sewer catchment to the monitoring data from AAW. The estimated and monitored loads were very similar (Table 6): 7.47 versus 5.30 kg day^−1^ for all PAOs and 7.79 versus 5.77 kg day^−1^ for all GAOs.

**Table 6.**
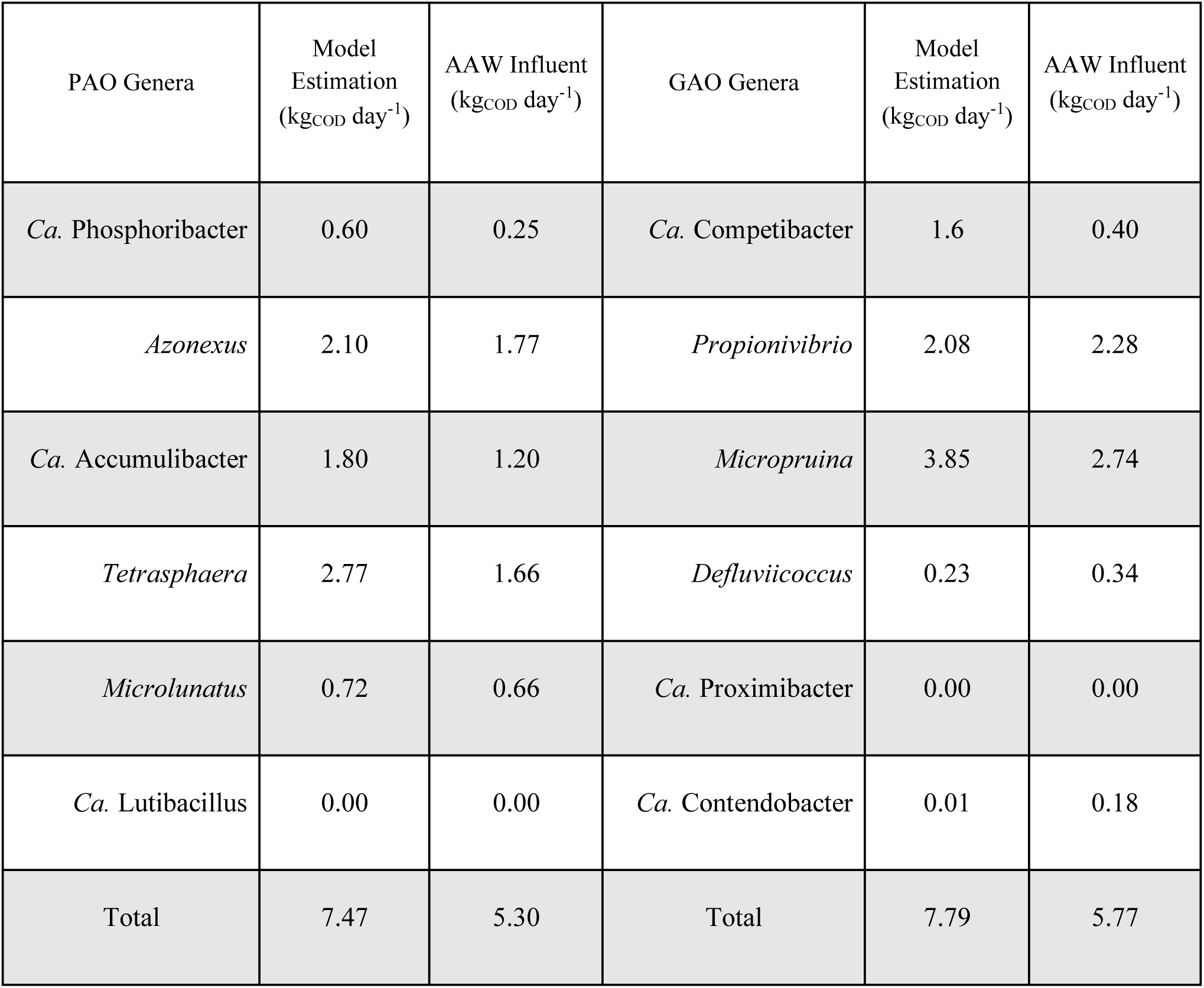
Comparison between PAOs and GAOs loads from the Aalborg sewer catchment to the AAW, estimated by the proposed sewer model and from the WWTP monitoring data.

Overall, the model estimates loads for all genera of PAO and GAO aligned very well with the loads estimated from AAW influent wastewater compositional data monitoring. For instance, the model-estimated load of *Azonexus*, the most abundant PAO in IWW to AAW was 2.10 kg day^−1^ compared to 1.77 kg day^−1^ from AAW influent wastewater monitoring. *Ca.* Accumulibacter had an estimated load of 1.80 kg day^−1^ compared to the monitored 1.20 kg day^−1^. However, the model overestimated some genera. For example, *Ca.* Phosphoribacter, the most abundant PAO in Danish AS systems (Dueholm et al., 2022), had an estimated load of 0.60 kg day^−1^ versus 0.25 kg day^−1^ from the monitoring data. *Ca.* Phosphoribacter is very low abundant in the AAW influent, making this genus more susceptible to detection uncertainties. Similarly, *Ca.* Competibacter had an estimated load of 1.62 kg day^−1^ compared to 0.40 kg day^−1^ from monitoring data. The presence of *Ca.* Competibacter shows strong seasonal patterns in AS, with recurring peaks between August and November in Danish WWTPs (Peces et al., 2022). Although it is unclear whether these patterns also occur in influent wastewater, this uncertainty likely contributes to the difficulty of accurately estimating *Ca.* Competibacter loads and other loads as the values shown are yearly averages. Further discrepancies between model-based and monitoring-based microbial abundances might be attributed to the nature of the sewer compositional data used in the model-based estimations. That dataset was based on grab samples taken from the Aalborg municipality sewer systems (Riisgaard-Jensen et al., 2025), which, at times, might fail to represent the complexity of the entire AAW sewer catchment.

The adequate performance of EBPR processes relies on proper operational conditions and the extent of bacterial immigration, which is known to influence the assembly of microbial communities in biological WWTPs, influencing processes such as phosphorus removal (Dottorini et al., 2021; Oh and Kim, 2021). Despite that, literature currently lacks efforts towards assessing bacterial loads from sewer catchments to WWTPs. In this sense, computational modeling can be utilized as a valuable tool for estimating biomass loads from sewer systems to WWTPs, helping to provide insights into microbial immigration without requiring extensive wastewater laboratory analyses. This contribution represents the first steps into this direction, offering a framework for further research aimed at improving current understanding on sewer microbial dynamics and potentially enhancing the design of sewer systems and WWTPs.

## 4. Conclusions

In this study, we developed a conceptual model to assess the biomass loads from gravity and pressure sewer systems, helping to shed light on the extent of which the sewer microbiome conveys bacteria towards WWTPs. We showed that gravity sewers produce higher overall biomass loads than pressure systems and that the microbial composition differs between the two systems. One of the features of our sewer model is its ability to estimate biomass loads from specific taxa. Overall, these estimations correlated well with figures calculated based on IWW data monitoring. For instance, the model-estimated loads for the PAOs *Azonexus*, the most abundant PAO in AAW’s influent wastewater, and *Ca.* Accumulibacter, the first identified and the most studied PAO, were 2.10 kg day^−1^ and 1.80 kg day^−1^, as compared to the IWW monitoring-based loads of 1.77 kg day^−1^ and 1.20 kg day^−1^, respectively. While providing a framework for further research on sewer microbial dynamics, this study offers a tool for gaining insights into microbial immigration without the need for extensive wastewater laboratory analyses and WWTPs monitoring, generating valuable information for improving microbial communities management in WWTPs, optimizing nutrient removal, and enhancing WWTP design. Future studies could try modeling transient hydraulic conditions to evaluate the impact of rain events on sewer biomass production.

## 5. Acknowledgements

This research was part of the RecaP project which has received funding from the European Union’s Horizon 2020 research and innovation programme under the Marie Skłodowska-Curie grant agreement No 956454.

