## Supplementary information for "Beyond Conveyance: Assessing the Contribution of Process-Critical Bacteria from Urban Sewer Systems to Wastewater Treatment Plants"

**Supplementary Material**

Table S1. Model parameters and initial variables definition.

| Symbol | Definition | Value | Unit | Reference |
| --- | --- | --- | --- | --- |
| S_F_ | Fermentable substrate | 18 | g_COD_ /m^3^ | Hvitved-Jacobsen et al., 2000 |
| S_A_ | Fermentation products | 18 | g_COD_ /m^3^ | ´´ |
| S_S_ | Readily biodegradable substrate (S_F_ + S_F_) | 36 | g_COD_ /m^3^ | ´´ |
| X_S1_ | Fast hydrolysable substrate | 90 | g_COD_ /m^3^ | ´´ |
| X_S2_ | Slow hydrolysable substrate | 444 | g_COD_ /m^3^ | ´´ |
| X_Bw_ | Biomass in the water phase | 30 | g_COD_ /m^3^ | Foladori et al., 2010 |
| COD | Total substrate | 600 | g_COD_ /m^3^ | Hvitved-Jacobsen et al., 2000 |
| µ_h_ | Aerobic heterotrophic biomass maximum growth rate | 3.25 | d^-1^ | Hvitved-Jacobsen et al., 1998 |
| K_S_ | Saturation constant for readily biodegradable substrate | 1.0 | g_COD_ /m^3^ | Hvitved-Jacobsen et al., 2000 |
| Y_h_ | Aerobic heterotrophic biomass yield constant | 0.55 | g_COD_/g_COD_ | ´´ |
| µ_fer_ | Fermentative biomass maximum growth rate | 3 | d^-1^ | ´´ |
| K_fer_ | Saturation constant of fermentable substrate for fermentative biomass | 20 | g_COD_ /m^3^ | Sun et al., 2014; Sun et al., 2018 |
| Y_fer_ | Fermentative biomass yield constant | 0.15 | g_COD_/g_COD_ | Gavala et al., 2003 |
| µ_ac_ | Acetogenic biomass maximum growth rate | 3 | d^-1^ | Sun et al., 2014; Sun et al., 2018 |
| K_ac_ | Saturation constant of fermentation products for acetogenic biomass | 500 | g_COD_ /m^3^ | ´´ |
| Y_ac_ | Acetogenic biomass yield constant | 0.15 | g_COD_/g_COD_ | ´´ |
| µ_met_ | Methanogenic biomass maximum growth rate | 0.24 | d^-1^ | ´´ |
| K_met_ | Saturation constant of fermentation products for methanogenic biomass | 409 | g_COD_ /m^3^ | ´´ |
| Y_met_ | Methanogenic biomass yield constant | 0.05 | g_COD_/g_COD_ | ´´ |
| µ_sul_ | Sulfidogenic biomass maximum growth rate | 0.55 | d^-1^ | ´´ |
| K_sul_ | Saturation constant of fermentation products for sulfidogenic biomass | 4.10 | g_COD_ /m^3^ | ´´ |
| Y_sul_ | Sulfidogenic biomass yield constant | 0.057 | g_COD_/g_COD_ | ´´ |
| K_O_ | Saturation constant for dissolved oxygen | 0.5 | g_O2_/ m^3^ | Hvitved-Jacobsen et al., 1998 |
| q_m_ | Maintenance constant for aerobic heterotrophs | 1.0 | d^-1^ | Hvitved-Jacobsen et al., 2000 |
| k_dec,ac_ | Decay rate of acetogenic biomass | 0.02 | d^-1^ | Sun et al., 2014; Sun et al., 2018 |
| k_dec,met_ | Decay rate of methanogenic biomass | 0.02 | d^-1^ | ´´ |
| k_dec,sul_ | Decay rate of sulfidogenic biomass | 0.02 | d^-1^ | ´´ |
| k_h1_ | Fast hydrolysis constant rate | 4.0 | d^-1^ | Hvitved-Jacobsen et al., 1998 |
| k_h2_ | Slow hydrolysis constant rate constant | 1.0 | d^-1^ | ´´ |
| K_X1_ | Saturation constant for fast hydrolysis | 0.5 | g_COD_/g_COD_ | ´´ |
| K_X2_ | Saturation constant for slow hydrolysis | 0.2 | g_COD_/g_COD_ | ´´ |
| h_A_ | Anaerobic hydrolysis reduction factor | 0.2 | - | Hvitved-Jacobsen et al., 2000 |
| ɛ | Efficiency constant for biofilm biomass | 0.15 | - | ´´ |
| X_Bf_ | Active biomass in the biofilm | 10 | g_COD_ /m^2^ | ´´ |
| ᵹ_anaerob_ | Anaerobic biofilm thickness | 0.6 | mm | Hvitved-Jacobsen et al., 2013; Sun et al., 2014 |
| ᵹ_aerob_ | Aerobic biofilm thickness | 1 | mm | Hvitved-Jacobsen et al., 2013,Li et al., 2019 |
| ɵ_w_ | Temperature coefficient water phase | 1.07 | - | Hvitved-Jacobsen et al., 2000 |
| ɵ_bio_ | Temperature coefficient biofilm | 1.05 | - | ´´ |
| ɵ_r_ | Temperature coefficient reaeration | 1.024 | - | ´´ |

Reaeration, in gravity sewers, was calculated considering the expression shown in Equation S1 (Jensen, 1995; Tanaka and Hvitved-Jacobsen, 2002) for estimating the overall oxygen transfer coefficient:

$r_{DO}=K_{La}\left( S_{O_{S}}-S_{O} \right)$, $K_{La}=0.86(1+0.2{F_{r}}^{2})(s \cdot v)^{\frac{3}{8}}{d_{m}}^{-1}$	(S1)

where:

$r_{DO}$ is the reaeration rate (g m^-3^h^-1^);

$K_{La}$ is the oxygen transfer coefficient at 20 $℃$ (h^-1^);

$S_{O_{S}}$ is the saturation dissolved oxygen concentration in wastewater at 20 (gO_2_ m^-3^);

$S_{O}$ is the dissolved oxygen concentration (gO_2_ m^-3^);

$F_{r}$ is the Froude number (-);

$s$ is the slope (m m^-1^);

$v$ is the mean flow velocity (m s^-1^);

$d_{m}$ is the hydraulic mean depth (m).

The temperature effect on reaction and reaeration rates was considered using the expression shown in Equation S2 (Tanaka and Hvitved-Jacobsen, 2002):

$r=r_{20}{\theta_{T}}^{\left( T-20 \right)}$	(S2)

where:

T is the temperature ($℃$);

$r$ is reaction rate of a concerned biochemical process at temperature T (g m^-3^h^-1^);

$r_{20}$ is the reaction rate of the concerned process at 20 $℃$ (g m^-3^h^-1^);

$\theta_{T}$ is a coefficient describing temperature dependency.

Table S2. Kinetic expressions for gravity sewer microbial transformations.

| Heterotrophic biomass growth, water | $r_{growthW}=\mu_{h}\frac{S_{S}}{K_{S}+S_{S}}\frac{S_{O}}{K_{O}+S_{O}}$ |
| --- | --- |
| Heterotrophic biomass growth, biofilm | $r_{growthB}=r_{growthW}ɛ$ |
| Maintenance energy requirement for heterotrophic biomass, water | $r_{maintenanceW}=q_{m}\frac{S_{O}}{K_{O}+S_{O}}$ |
| Maintenance energy requirement for heterotrophic biomass, biofilm | $r_{maintenanceB}=r_{maintenanceW}ɛ$ |
| Hydrolysis – fast hydrolysable substrate, water | $r_{S1w}=k_{h1}\frac{\frac{X_{S1}}{X_{W}}}{K_{X1}+\frac{X_{S1}}{X_{W}}}$ |
| Hydrolysis – fast hydrolysable substrate, biofilm | $r_{S1b}=r_{S1w} ɛ$ |
| Hydrolysis – slow hydrolysable substrate, water | $r_{S2w}=k_{h2}\frac{\frac{X_{S2}}{X_{W}}}{K_{X2}+\frac{X_{S2}}{X_{W}}}$ |
| Hydrolysis – slow hydrolysable substrate, biofilm | $r_{S2b}=r_{S2w} ɛ$ |

Table S3. Gravity sewers model matrix.

|  | $S_{S}$ | $X_{S1}$ | $X_{S2}$ | $X_{w}$ | $S_{O}$ | rate |
| --- | --- | --- | --- | --- | --- | --- |
| Suspended biomass growth | -1/ Y_h_ |  |  | 1 | -(1-Y_h_)/ Y_h_ | r_growthW_ |
| Biofilm growth | -1/ Y_h_ |  |  | 1 | -(1-Y_h_)/ Y_h_ | r_growthB_ |
| Maintenance energy requirement – suspended biomass | -1 |  |  |  | -1 | r_maintenanceW_ |
| Maintenance energy requirement – biofilm | -1 |  |  |  | -1 | r_maintenanceB_ |
| Suspended hydrolysis, fast | 1 | -1 |  |  |  | r_S1w_ |
| Biofilm hydrolysis, fast | 1 | -1 |  |  |  | r_S1b_ |
| Suspended hydrolysis, slow | 1 |  | -1 |  |  | r_S2w_ |
| Biofilm hydrolysis, slow | 1 |  | -1 |  |  | r_S2b_ |
| Reaeration |  |  |  |  | 1 | r_DO_ |

For the pressure sewers modeling, as a simplification, the microbial biomass in pressure system biofilms was assumed to be composed of acetogenic, methanogenic, and sulfidogenic microorganisms in proportions of 50%, 44%, and 6%, based on the findings of Hvitved-Jacobsen *et al.* (2013) and Sun *et al.* (2014). Hence, the pressure sewers biofilm biomass maximum growth rate was calculated as described in Equation S3, the pressure sewers biofilm biomass saturation constant for fermentation products was calculated as described in Equation S4, and the pressure sewers biofilm biomass yield constant was calculated as described in Equation S5.

${\mu_{Abio}={(0.5\mu}_{ac}}{+0.44\mu}_{met}{+0.06\mu}_{sul})$	(S3)

${K_{Abio}= (0.5K_{ac}}{+0.44K}_{met}{+0.06K}_{sul})$	(S4)

$Y_{Abio}= {(0.5Y}_{ac}{+0.44Y}_{met}{+0.06Y}_{sul})$	(S5)

Table S4. Kinetic expressions for pressure sewer microbial transformations.

| Fermentative biomass growth, water | $r_{AgrowthW}=\mu_{fer}\frac{S_{F}}{K_{fer}+S_{F}}$ |
| --- | --- |
| Acetogenenic, methanogenenic, and sulfidogenenic biomass growth, biofilm | $r_{AgrowthB}=\mu_{Abio}\frac{S_{A}}{K_{Abio}+S_{A}}ɛ$ |
| Maintenance energy requirement for fermentative biomass | $r_{AmaintenanceW}=k_{dec,ac}=k_{dec,met}{=k}_{dec,sul}$ |
| Maintenance energy requirement for biofilm biomass | $r_{AmaintenanceB}=r_{AmaintenanceW}ɛ$ |
| Anaerobic hydrolysis – fast hydrolysable substrate, water | $r_{AS1}=k_{h1}\frac{\frac{X_{S1}}{X_{W}}}{K_{X2}+\frac{X_{S2}}{X_{W}}}h_{A}$ |
| Anaerobic hydrolysis – fast hydrolysable substrate, biofilm | $r_{AS1b}=r_{AS1}ɛ$ |
| Anaerobic hydrolysis – slow hydrolysable substrate, water | $r_{AS2}=k_{h2}\frac{\frac{X_{S2}}{X_{W}}}{K_{X2}+\frac{X_{S2}}{X_{W}}}h_{A}$ |
| Anaerobic hydrolysis – slow hydrolysable substrate, biofilm | $r_{AS2b}=r_{AS2}ɛ$ |

Table S5. Pressure sewers model matrix.

|  | $S_{F}$ | $S_{A}$ | $X_{S1}$ | $X_{S2}$ | $X_{w}$ | rate |
| --- | --- | --- | --- | --- | --- | --- |
| Fermentation – water | -1/ Y_fer_ | 1/ Y_fer_ |  |  | 1 | r_AgrowthW_ |
| Acetogenesis, methanogenesis, sulfidogenesis – biofilm |  | -1/ Y_Abio_ |  |  | 1 | r_AgrowthB_ |
| Anaerobic hydrolysis, fast – water | 1 |  | -1 |  |  | r_AS1_ |
| Anaerobic hydrolysis, fast – biofilm | 1 |  | -1 |  |  | r_AS1b_ |
| Anaerobic hydrolysis, slow – water | 1 |  |  | -1 |  | r _AS2_ |
| Anaerobic hydrolysis, slow – biofilm | 1 |  |  | -1 |  | r _AS2b_ |
| Maintenance energy requirement – suspended biomass | -1 |  |  |  |  | r_AmaintenanceW_ |
| Maintenance energy requirement – biofilm |  | -1 |  |  |  | r_AmaintenanceB_ |

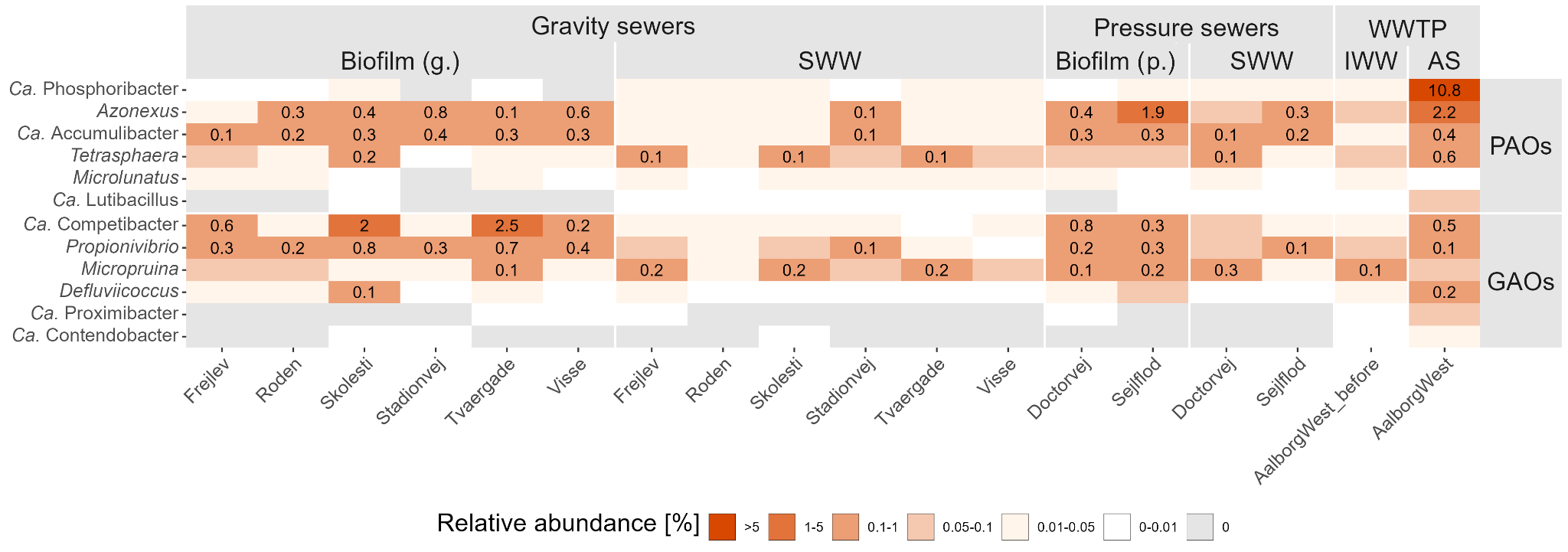

Figure S1. Mean relative abundance of polyphosphate accumulating bacteria (PAO) and glycogen accumulating bacteria (GAO) across all the sampling locations. The facets show whether the genera are a PAO or GAO. Abundances are shown with two decimals if ≥0.01. A relative abundance of 0 means no reads were detected.

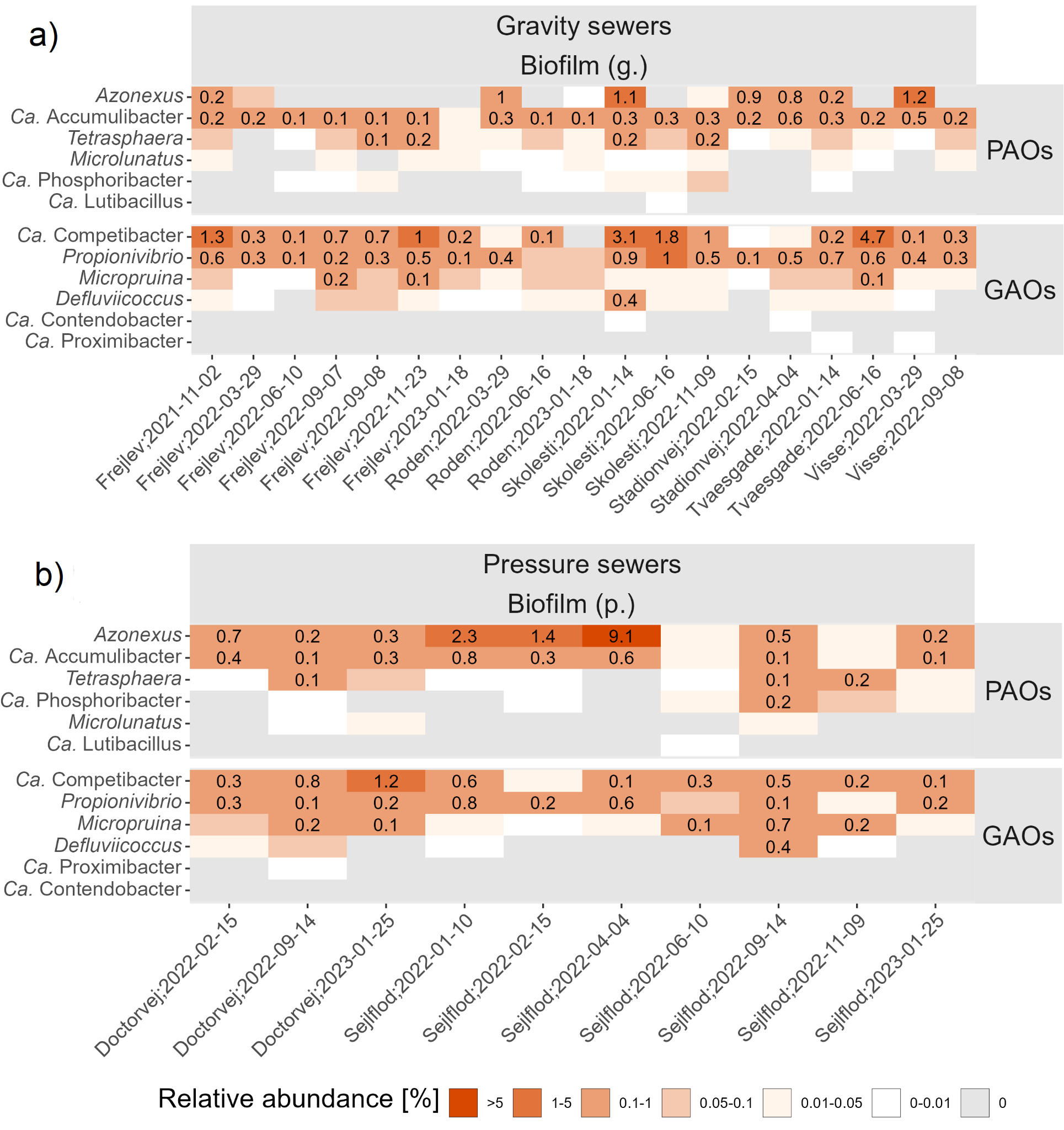

Figure S2. Mean relative abundance of polyphosphate accumulating bacteria (PAO) and glycogen accumulating bacteria (GAO) across all biofilm samples collected a) in gravity sewers, and b) in pressure sewers. The facets show whether the genera are a PAO or GAO. Abundances are shown with two decimals if ≥0.01. A relative abundance of 0 means no reads were detected.

**Supplementary Material - References**

Foladori, P., Bruni, L., Tamburini, S., & Ziglio, G. (2010). Direct quantification of bacterial biomass in influent, effluent and activated sludge of wastewater treatment plants by using flow cytometry. Water Research, 44(13), 3807-3818. https://doi.org/10.1016/j.watres.2010.04.027

Gavala, H. N., Angelidaki, I., & Ahring, B. K. (2003). Kinetics and modeling of anaerobic digestion process. Biomethanation I, 57-93. https://doi.org/10.1007/3-540-45839-5_3

Hvitved-Jacobsen, T., Vollertsen, J., & Nielsen, A. H. (2013). Sewer processes: microbial and chemical process engineering of sewer networks (2nd ed.). CRC Press.

Hvitved-Jacobsen, T., Vollertsen, J., & Nielsen, P. H. (1998). A process and model concept for microbial wastewater transformations in gravity sewers. Water Science and Technology, 37(1), 233-241. https://doi.org/10.1016/S0273-1223(97)00774-9

Hvitved-Jacobsen, T., Vollertsen, J., & Tanaka, N. (2000). An integrated aerobic/anaerobic approach for prediction of sulfide formation in sewers. Water Science and Technology, 41(6), 107-115. https://doi.org/10.2166/wst.2000.0099

Jensen, N. A. (1995). Empirical modeling of air‐to‐water oxygen transfer in gravity sewers. Water Environment Research, 67(6), 979-991. https://doi.org/10.2175/106143095X133211

Li, W., Zheng, T., Ma, Y., & Liu, J. (2019). Current status and future prospects of sewer biofilms: Their structure, influencing factors, and substance transformations. Science of the Total Environment, 695, 133815. https://doi.org/10.1016/j.scitotenv.2019.133815

Sun, C., Zhang, B., Ning, D., Zhang, Y., Dai, T., Wu, L., ... & Wen, X. (2021). Seasonal dynamics of the microbial community in two full-scale wastewater treatment plants: diversity, composition, phylogenetic group based assembly and co-occurrence pattern. Water Research, 200, 117295. https://doi.org/10.1016/j.watres.2021.117295

Sun, J., Hu, S., Sharma, K. R., Ni, B. J., & Yuan, Z. (2014). Stratified microbial structure and activity in sulfide-and methane-producing anaerobic sewer biofilms. Applied and Environmental Microbiology, 80(22), 7042-7052. https://doi.org/10.1128/AEM.02146-14

Tanaka, N., & Hvitved-Jacobsen, T. (2002). Anaerobic transformations of wastewater organic matter and sulfide production–investigations in a pilot plant pressure sewer. Water science and Technology, 45(3), 71-79. https://doi.org/10.2166/wst.2002.0057
